# Selective Agonism of Liver and Gut FXR Prevents Cholestasis and Intestinal Atrophy in Parenterally Fed Neonatal Pigs

**DOI:** 10.1101/2024.09.03.611073

**Authors:** Yanjun Jiang, Zhengfeng Fang, Gregory Guthrie, Barbara Stoll, Shaji Chacko, Sen Lin, Bolette Hartmann, Jens J. Holst, Harry Dawson, Jose J. Pastor, Ignacio R. Ipharraguerre, Douglas G. Burrin

## Abstract

**BACKGROUND & AIMS:** We aimed to investigate the relative efficacy of feeding different bile acids in preventing PNALD in neonatal pigs.

**METHODS:** Newborn pigs given total parenteral nutrition (TPN) combined with minimal enteral feeding of chenodeoxycholic acid (CDCA), or increasing doses of obeticholic acid (OCA) for 19 days.

**RESULTS:** Enteral OCA (5 and 15 mg/kg), but not CDCA (30 mg/kg) reduced blood cholestasis markers compared to TPN controls and increased bile acids in the gallbladder and intestine. Major bile acids in the liver and distal intestine were CDCA, HCA, HDCA and OCA, and their relative proportions were increased by the type of bile acid (CDCA or OCA) given enterally. High doses of OCA increased the total NR1H4-agonistic bile acid profile in the liver and intestine above 50% total bile acids. Both CDCA and OCA treatments suppressed hepatic cyp7a1 expression, but only OCA increased hepatobiliary transporters, ABCB11, ABCC$ and ABCB1. Plasma phytosterol levels were reduced and biliary levels were increased by CDCA and OCA and hepatic sterol transporters, abcg5/8, expression were increased by OCA. Both CDCA and OCA increased plasma FGF19 and OCA increased intestinal FGF19, FABP6, and SLC51A. Both CDCA and OCA increased intestinal mucosal growth, whereas CDCA increased the plasma GLP-2, GLP-1 and GIP.

**CONCLUSIONS:** Enteral OCA prevented cholestasis and phytosterolemia by increased hepatic bile acid and sterol transport via induction of hepatobiliary transporter FXR target genes and not by suppression of bile acid synthesis genes. We also showed an intestinal trophic action of OCA that demonstrates a dual clinical benefit of FXR agonism in the prevention of PNALD in piglets.

## Introduction

Parenteral nutrition (PN) is a life-saving means of nutritional support for thousands of hospitalized infants that cannot tolerate enteral nutrition due to the immaturity or surgical removal of the intestine due to congenital or disease-related causes. Administration of PN increases the risk for metabolic condition often termed parenteral nutrition-associated liver disease (PNALD) or intestinal failure-associated liver disease (IFALD) in cases where much of the intestine has been surgically resected^1, 2^. The incidence of PNALD can be as high as 50% in premature infants given total PN (TPN) for prolonged periods and is marked by increased serum concentrations of conjugated bilirubin and bile acids along with hepatic steatosis, biliary ductal inflammation and eventually fibrosis ^3^. Current clinical strategies to prevent or treat PNALD in infants include various PN lipid emulsion approaches and enteral ursodeoxycholic acid (UDCA) treatment ^4–6^.

A molecular mechanism implicated in the cause of PNALD is that plant phytosterols, enriched in soybean-oil lipid emulsions, disrupt the function of the liver farnesoid X receptor (NR1H4), which is the primary sensor of bile acids that controls the molecular regulation of targets genes involved in bile acid homeostasis. Reports in cultured pig, mice and human hepatocytes show that phytosterols antagonize FXR. Studies in PN-fed mice showed that addition of the phytosterol stigmasterol to fish oil emulsion induced PNALD but only in the presence of intestinal injury and inflammation^7–9^. In contrast, our study in TPN-fed piglets showed that addition of phytosterols to fish oil emulsion did not induce PNALD^10^. Fibroblast growth factor 19 (FGF19) is an enterokine hormone released from the distal intestine in response to luminal bile acid activation of FXR in enterocytes and it functions as a negative feedback signal for hepatic CYP7A1 mediated bile synthesis and promotes gallbladder filling^11–13^.

Parenteral nutrition-induced cholestasis disrupts bile acid-mediated activation of intestinal FXR-FGF19 signaling and we showed that treatment with enteral chenodeoxycholic acid (CDCA) restored FGF19 secretion and reduced serum cholestasis in neonatal pigs^14^. Enteral CDCA treatment of PN-fed pigs also restored intestinal mucosal growth and secretion of glucagon-like peptide 2 (GLP-2) implicating activation of enteroendocrine GPBAR1 receptors^14^. A selective FXR agonist obeticholic acid (OCA) is approved for use in treatment of primary biliary cholangitis in adults^15^. Selective activation of hepatic FXR with OCA in PN-fed mice prevented PNALD by restoration of hepatic NR1H4 gene expression of canalicular bile and of sterol and phospholipid transporters and suppression of Kupffer cell activation^16^. Here we show that selective agonism of both liver and intestine FXR using an enteral strategy prevents cholestasis and intestinal atrophy in neonatal PN-fed pigs. Our findings demonstrate a therapeutic approach to restore gut-liver bile acid signaling with clinically available FXR agonist in the prevention of intestinal atrophy and PNALD in PN-fed infants.

## Materials and Methods

### Animals and surgery

Neonatal domestic crossbred pigs were obtained from a commercial swine farm and housed in the Children’s Nutrition Research Center. Piglets (2-d-old) were surgically implanted with catheters in the jugular vein and duodenum as described previously^14^. Piglets were administered intramuscular injections of iron dextran (100 mg), ampicillin (50 mg/kg), and buprenorphine (0.01 mg/kg) prior to surgery; ampicillin (50 mg/kg) was given daily thereafter until the end of the study. Piglets received TPN administered via implanted jugular catheters as described previously ^14, 17^. The study protocol was approved by the Animal Care and Use Committee of Baylor College of Medicine and was conducted in accordance with the Guide for the Care and Use of Laboratory Animals (DRR/NIH, Bethesda, MD).

### Nutritional support, bile acid treatment and study design

Total parenteral nutrition consisted of a solution containing a complete nutrient mixture of amino acids, glucose, electrolytes, vitamins, and trace minerals, administered via jugular catheter, and a parenteral lipid emulsion (10 g/kg/day; 20% Intralipid). All TPN piglets received the following amounts of nutrition per kilogram body weight: fluid, 265 ml; energy, 246 kcal; carbohydrate, 25g; protein, 14 g; and lipid, 10 g, as described ^17–19^. Post-surgery, TPN was started at 5 ml/(kgꞏh) and gradually increased to 11 ml/(kgꞏh). The bile acids used were solubilized in ethanol in order to be infused. Pigs were randomly assigned to receive intraduodenal total daily infusion of either sham (ethanol, 0.15 ml/kg), chenodeoxycholic acid (CDCA) (30 mg/kg) or one of three obeticholic acid (OCA) doses (0.5, 5, 15 mg/kg) divided into three times per day at 0700 h, 1500 h, and 2300 h for 19 days. On day 19, jugular blood was collected and pigs were given an intravenous injection of 5-bromodeoxyuridine (BrdU, 50 mg/kg; Sigma Aldrich, St. Louis, MO) 4 h before being euthanized with an intravenous injection of pentobarbital sodium (50 mg/kg) and phenytoin sodium (5 mg/kg; Beutanasia-D; Schering-Plough Animal Health, Kenilworth, NJ). Total contents of the gallbladder were collected by needle aspiration and livers were dissected and weighed. Luminal contents from the small intestine and proximal colon were sampled by flushing with sterile saline. Liver and intestinal tissues were isolated, weighed and samples were frozen in liquid nitrogen and stored at −80°C until analysis. Liver and intestinal samples also were fixed in OCT and frozen at −80°C and as well as in 10% paraformaldehyde for 24 hours, then transferred to 70% ethanol before paraffin embedding and stored for later analysis.

### Blood and tissue analysis

Blood samples were collected in Na2EDTA tubes and serum tubes and processed to obtain plasma and serum, respectively. Plasma and serum samples were stored at −80°C until further analyses. Blood samples also were collected and serum chemistry markers were analyzed on a Roche-Cobas 6000 analyzer as described previously^10^. Plasma FGF19 was measured using the porcine FGF-19 ELISA kit (RayBiotech). Plasma glucagon like peptide 1 and 2 (GLP-1 and GLP-2) and glucose-dependent insulinotropic polypeptide (GIP) were assayed as described previously ^14, 19, 20^. Intestinal samples were stained with hematoxylin and eosin and villus height and crypt depth were measured using an Axiophot microscope (Carl Zeiss) and Image version 1.60 software (National Institutes of Health) as described previously^21^. Crypt cell proliferation was measured by quantifying BrdU positive crypt cells as described previously^21^. Immunohistochemical pancytokeratin (pan CK) staining was performed on paraffin-embedded liver sections using standard protocols. The pan CK antibody (monoclonal mouse anti-human, DAKO, Agilent, M3515) was used at a dilution of 1:500. Prior to antibody incubation, antigen retrieval was performed with citrate buffer for improved binding. Slides were stained with chromogen diaminobenzidine (DAB) substrate for visualization. Counterstain with hematoxylin was used to visualize nuclei. Stained slides were scanned using a Zeiss Axio scan.Z1 (Zeiss, Gottingen, Germany) slide scanner. From the full tissue image, 6 randomly selected fields of equal size were converted into image files for downstream stain quantification. ImageJ was used for quantification as follows per field image. Image colors were separated using color deconvolution with H-DAB setting to get both DAB staining (Pan CK positive cells) and Hematoxylin staining (nuclei). Images were converted to binary, then watershed separation was applied to identify individual cells that were clustered. The cell numbers were then counted using the analyze particles function with a minimum particle size of 100 pixels set to remove background noise. The number of cholangiocytes was then normalized to the number of nuclei per field.

### Bile Acid and Phytosterol Analysis

Total bile acid concentrations were measured in plasma, gallbladder bile and tissue using an enzymatic colorimetric kit as described previously^22^. For liver, bile, and small intestine the bile acid concentration was measured in supernatant after homogenization in ethanol. Total bile acid pool size of liver tissue, gallbladder, and intestine tissue was calculated as the product of the total bile acid concentration (unit/g tissue) and the mass of the organ or compartment (g/kg BW) corrected for body weight at the end of study. Quantitation of bile acid profiles in liver and intestinal tissue was performed using ultra performance liquid chromatography (UPLC)-MS analysis of tissue extracted with 1:1 H_2_O-acetonitrile (ACN) containing chenodeoxycholic acid-d4 as internal standard as described previously^21, 23^. Bile acid pool hydrophobicity was estimated by multiplying bile acid species fractions by hydrophobicity indices estimated by Heuman ^24^. Quantitation of phytosterols (β-sitosterol, stigmasterol, campesterol) in the plasma and liver was performed using liquid chromatography tandem mass spectrometry (LC-MS/MS) as described previously^22^.

### Primary hepatocyte culture

Hepatocytes were isolated from three piglets (2 day old) using a previously described protocol ^25^. Hepatocytes were plated in 24-well plates at 5.0 x 10^5^ cells/well in William’ E media containing fetal bovine serum, glucagon, insulin-transferrin-selenium, and L-glutamine for 16 hours. Subsequently, media was changed to Williams E media without supplements for 8 hours. Hepatocytes were then treated with individual bile acids or combinations of bile acids to reflect the bile acid composition found within the piglets receiving TPN treatments in the current study. The treatment groups and their bile acid compositions were as follows: CON (DMSO 0.1%), HCA (50 µM hyocholic acid), HDCA (50 µM hyodeoxycholic acid), UDCA (50 µM ursodeoxycholic acid), CDCA (50 µM chenodeoxycholic acid), OCA (5 µM obeticholic acid), TPN-CON (31 µM hyocholic acid, 10 µM hyodeoxycholic acid, 9 µM chenodeoxycholic acid), TPN-CDCA (72 µM chenodeoxycholic acid, 11 µM hyodeoxycholic acid, 9.5 µM hyocholic acid, 1.5 µM ursodeoxycholic acid), TPN-OCA (22.5 µM chenodeoxycholic acid, 17.5 µM hyocholic acid, 5 µM hyodeoxycholic acid, 2.5 µM ursodeoxycholic acid, 2.5 µM obeticholic acid). Twenty-four hours following treatment, cells were washed with cold PBS and RNA was extracted for downstream analysis.

### Immunoblotting and real-time PCR

Western immunoblotting was performed as described previously (7). Mouse monoclonal CYP7A1 antibody (clone 1D9) was purchased from Cosmo Bio Co., LTD. (Tokyo, Japan). Rabbit polyclonal FGF19 antibody (Prestige) was purchased from Sigma-Aldrich (St. Louis, MI). For analysis of CYP7A1 protein level in livers from different pigs, frozen liver tissues were homogenized with 1ml RIPA buffer (Boston BioProuducts) containing protease inhibitors. The homogenates were sonicated and centrifuged at 12,000 g for 15 minutes at 4°C. After measuring protein concentrations of the extracts, 50 µg of total protein was loaded to detect CYP7A1. For detecting circulating FGF19 protein level in jugular and portal venous plasma from various treatment groups, 10 µl of plasma was loaded in each lane.

Quantitative real-time PCR was performed on frozen liver samples. The cDNA was generated from RNA extracted from 100 to 150 mg of frozen liver tissue as described previously ^18^. Real-time qPCR was performed with Sybr green chemistry (Applied biosystems) on a Bio Rad CFX96. Primers were designed using software from NCBI Primer Blast based on the predicted porcine sequence available on ENSEMBL (**Supplemental Table 1**). Amplification efficiency was controlled by the use of an internal control (GAPDH or actin). Relative quantification of target mRNA expression was calculated and normalized to GAPDH or actin expression. All reactions were performed under the following thermal cycling conditions: 10 min at 95°C followed by 40 cycles of 95°C for 15 s and 60°C for 60 s. The 2^-ΔΔCT^ method was used to compare gene expression levels between samples, which were analyzed to determine the fold induction of mRNA expression.

### Statistics

Data are presented as mean ± SEM. Statistical analysis was performed using Prism Graphpad 9.2.0. Differences among the five groups were first analyzed using one way ANOVA, and post-hoc analysis was done using Tukey’s test as described in the figure legends. P values < 0.05 were considered significant.

## Results

### Enteral Obeticholic Acid Treatment Reduces Cholestasis and Promotes Intestinal Bile Flow

To determine the relative efficacy of different bile acids to modulate gut-liver FXR-FGF19 signaling, we measured serum markers of cholestasis and bile acid pools in TPN-fed pigs given enteral CDCA (30 mg/kg) or OCA (0.5, 5, or 15 mg/kg) for 19 days. Treatment with OCA at 5 and 15 mg/kg doses, but not CDCA or 0.5 mg/kg OCA, reduced serum direct bilirubin, GGT and total bile acid concentrations compared to control TPN alone (Figure 1A). Liver bile acid pool size was higher in CDCA than in any of the other groups, whereas the gallbladder and intestinal bile acid pool sizes were higher in pigs treated with CDCA and high dose of OCA (Figure 1B). Measurements of liver weight and relative bile duct density were not different among all groups (Figure 1C).

**Figure 1.**
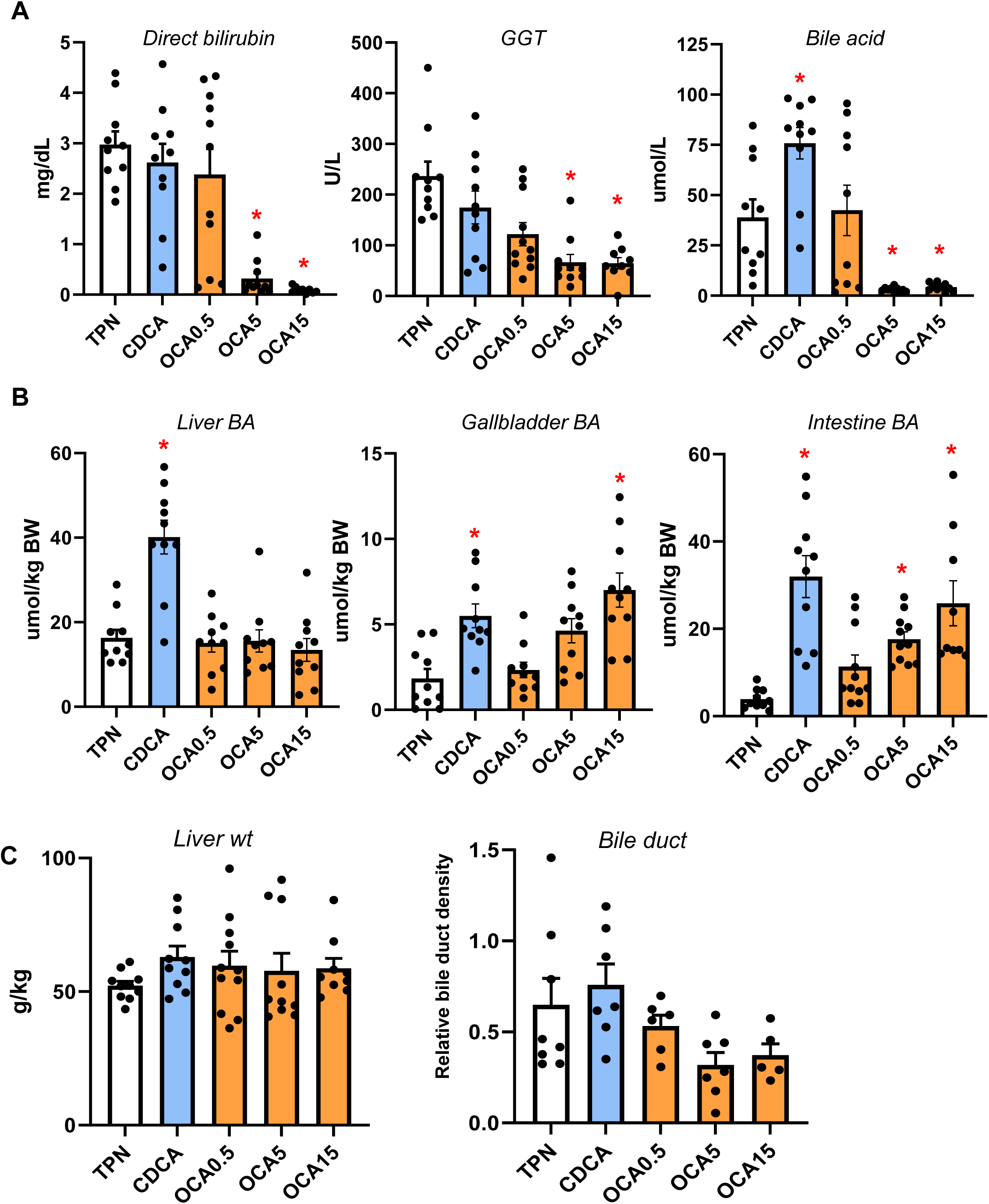
Serum and plasma markers of cholestasis and bile acid pool sizes of liver and intestine. Shown in top panel (A) are serum direct bilirubin and GGT and plasma total bile acid concentrations in treatment groups administered TPN, TPN+CDCA (30 mg/kg/d), TPN+OCA (0.5, 5.0, 15.0 mg/kg/d). Middle panel (B) shows total bile acid pool sizes for liver, gallbladder, and intestine. Bottom panel (C) shows liver weight and relative bile duct density measured as described in methods section.

### Enteral Obeticholic Acid Increases the Hepatic and Intestinal Bile Acid Profile Hydrophobicity and FXR Agonist Activity

We next measured the bile acid profiles of the most abundant natural bile acids in pigs CDCA, HCA, HDCA as well as OCA in the liver and distal small intestinal tissues (Figure 2). The data shown represent the relative amount total of conjugated and unconjugated bile acids in each tissue; approximately 98% of the bile acids were conjugated and 90-95% were conjugated to glycine compared to 5-10% taurine (not shown). The bile acid profiles in the liver and intestinal tissues were similar, which in control TPN pigs were dominated by HCA (60%), CDCA (20%), and HDCA (19%). In CDCA treated pigs, there was a significantly higher proportion of CDCA than control TPN pigs and CDCA (54%) was the dominant bile acid species followed by HCA (20%) and HDCA (22%). In control TPN pigs OCA was not detected, whereas its proportion in liver and intestine tissue increased as a function of the dose of OCA given to pigs 0.5 (9%), 5 (30%) and 15 (48%) mg/kg, respectively.

**Figure 2.**
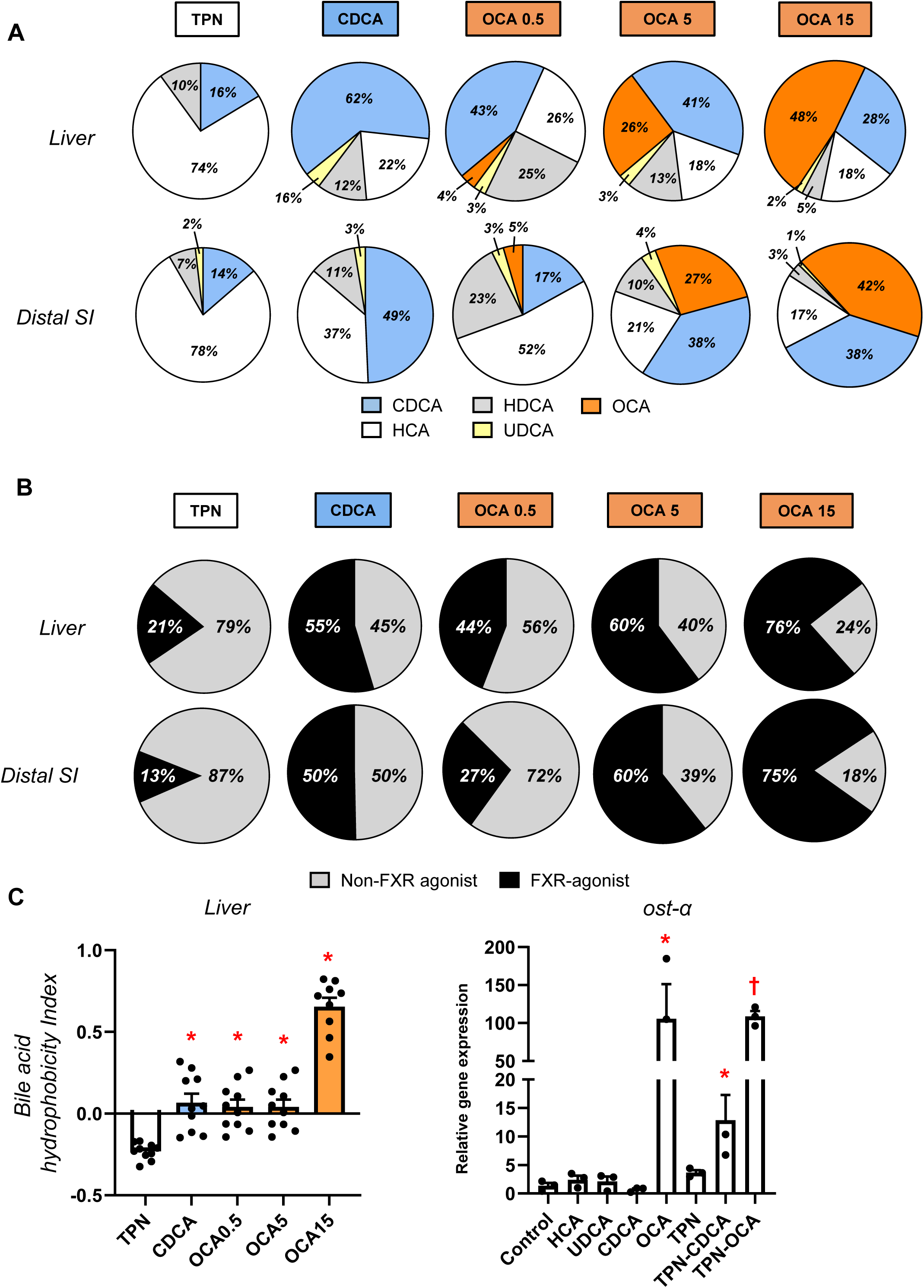
Liver and intestine bile acid profiles. Shown in top panel (A) are relative percentages of major bile acid species in the liver and distal small intestine (SI) in treatment groups administered TPN, TPN+CDCA (30 mg/kg/d), TPN+OCA (0.5, 5.0, 15.0 mg/kg/d). Middle panel (B) shows the estimated percentages of the bile acid pool with FXR-agonist and non-FXR agonist activity in the liver and distal SI tissue pools. Bottom panel (C) shows the calculated hydrophobicity index based on the measured liver tissue bile acid concentration as described in the methods section. Values are means with standard error, n=8-10 pigs/group. *P < 0.05 versus TPN, #P < 0.05. Also shown is the relative capacity of specific bile acids and mixtures of bile acids to induce SLC51A expression in cultured primary pig hepatocytes after 24 hr incubation. We modeled bile acid profiles based on the measured concentrations in liver tissue of TPN, TPN+CDCA, and TPN+OCA groups. Values represent mean with standard deviation of three separate replicates of each treatment. *P < 0.05 versus control, #P < 0.05.

Interestingly, the proportion of CDCA (25-35%) also tended to increase in the liver and intestine of OCA treated pigs compared to control TPN pigs. We also calculated the proportion of tissue bile acids that show FXR-agonist activity based on activation of the FXR-target gene SLC51A in cultured pig hepatocytes as well as their combined hydrophobicity (Figure 2 B-C) using an approach described previously ^26^. We found that the hydrophobicity of the bile acid pools of the CDCA and OCA treated groups were higher than in control TPN pigs. Pig hepatocyte cultures showed that CDCA and OCA hade potent FXR-agonist activity, whereas HCA and HDCA do not (Figure 2 B). Importantly, this was also reflected in the FXR-agonist activity of bile mixtures representing TPN vs CDCA and OCA treated pigs. These difference in FXR-agonist activity were translated into a progressively higher FXR-agonist profile in liver and intestinal tissues (Figure 2 C). It was remarkable that in both liver and intestinal tissue approximately 13-20% of the bile acids were FXR agonist in control TPN pigs, whereas the contribution of FXR-agonist species activity represented 50% in CDCA pigs and 60-75% in the two highest dose OCA groups.

### Enteral Obeticholic Acid Treatment Reduces Hepatic CYP7A1 Expression and Increased hepatobiliary Bile Acid Transporter Expression

To examine how the changes in bile acid profiles affected liver bile acid homeostasis, we measured the expression of various genes and protein involved in bile acid synthesis (Figure 3) and transport (Figure 4). The expression of hepatic CYP7A1 mRNA and protein abundance was suppressed by CDCA and OCA at tested doses. The expression of hepatic FXR target genes, NR0B2 and FGF19 was upregulated by the highest OCA dose, while genes involved in hepatobiliary bile acid transport, including ABCB11, ABCC4, SLC51A, and ABCB1, were increased by the intermediate and high OCA doses.

**Figure 3.**
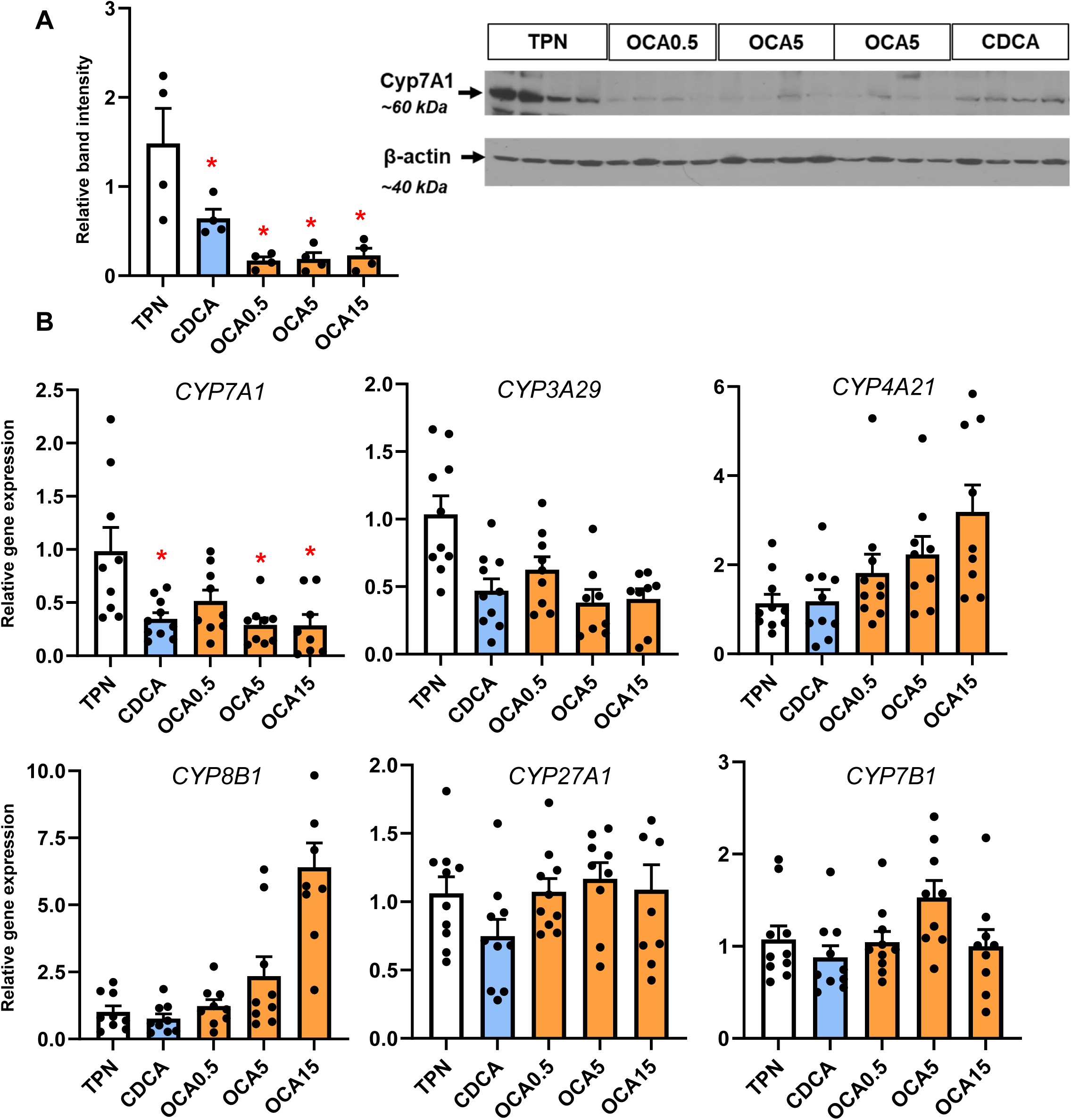
Hepatic expression of bile acid synthesis genes. Shown in top panel (A) is the expression of hepatic tissue cyp7A1 abundance by Western immunoblot in treatment groups administered TPN, TPN+CDCA (30 mg/kg/d), TPN+OCA (0.5, 5.0, 15.0 mg/kg/d). Botton panel (B) shows the expression of genes in the bile acid synthesis. Values are means with standard error, n=8-10 pigs/group. *P < 0.05 versus TPN, #P < 0.05.

**Figure 4.**
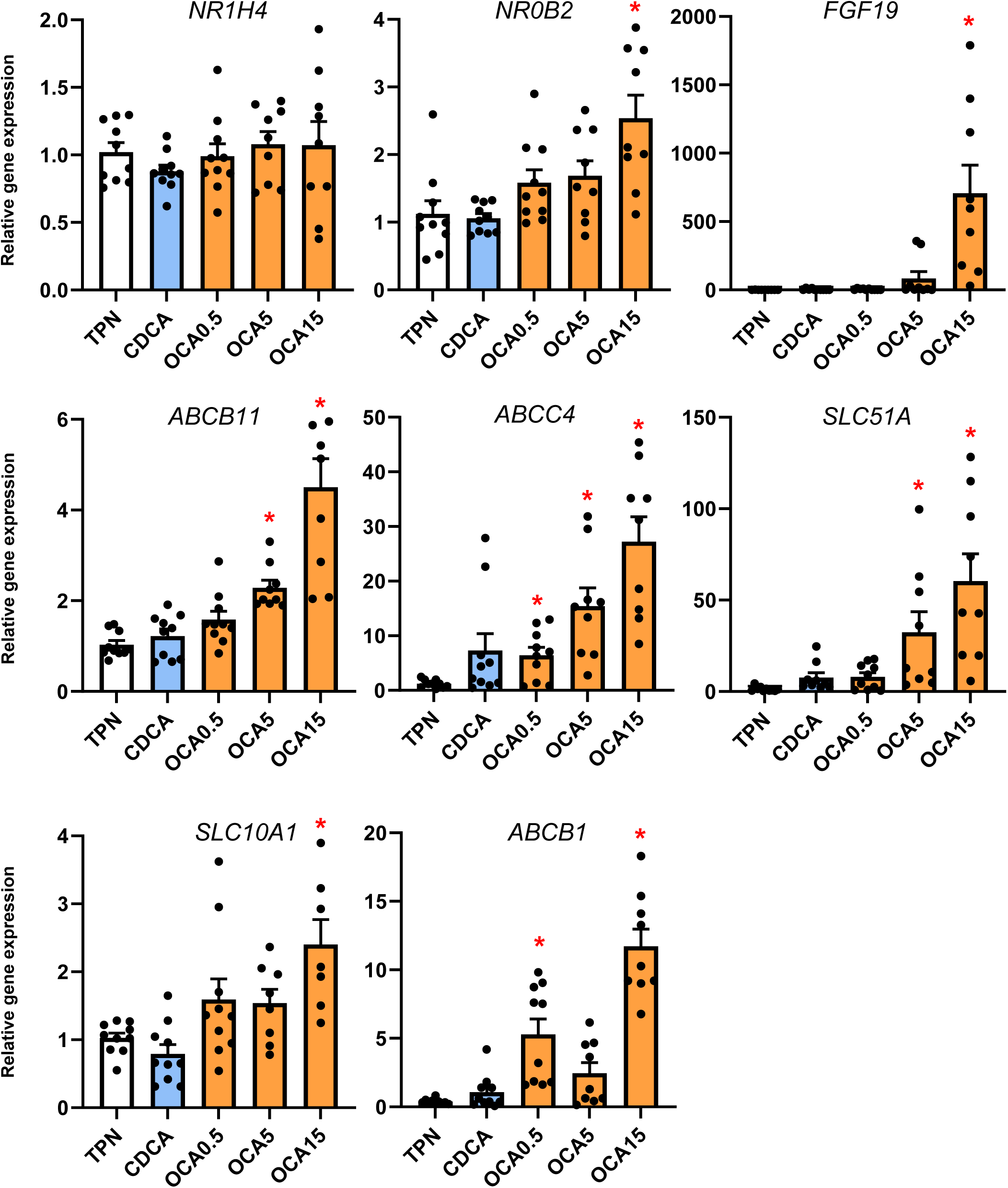
Liver FXR target gene and bile acid transport gene expression. Shown is the expression of hepatic tissue genes involved in bile acid transport in treatment groups administered TPN, TPN+CDCA (30 mg/kg/d), TPN+OCA (0.5, 5.0, 15.0 mg/kg/d). Values are means with standard error, n=8-10 pigs/group. *P < 0.05 versus TPN, #P < 0.05.

### Enteral Obeticholic Acid Treatment Reduces Plasma Phytosterolemia by Increasing Hepatobiliary Phytosterol Transport

Our previous studies showed that neonatal pigs given TPN including soybean oil-based lipid emulsions led to phytosterolemia secondary to cholestasis and blockage of hepatobiliary sterol transport ^18, 22^. We measured the concentration of phytosterols and cholesterol in plasma, liver and bile to assess whether changes in hepatic bile flow in response to enteral bile acid treatments altered liver sterol homeostasis (Figure 5). Enteral treatment with CDCA and all doses of OCA reduced plasma phytosterolemia, increased hepatic phytosterol content and biliary concentration consistent with increased biliary flow of bile acids (Figure 5 A). There were no differences in plasma total cholesterol concentration among treatment groups, but both CDCA and OCA treatments increased the biliary cholesterol content (Figure 5 B). Enteral OCA treatment also increased the hepatic expression of both sterol transporters, ABCG5 and ABCG8.

**Figure 5.**
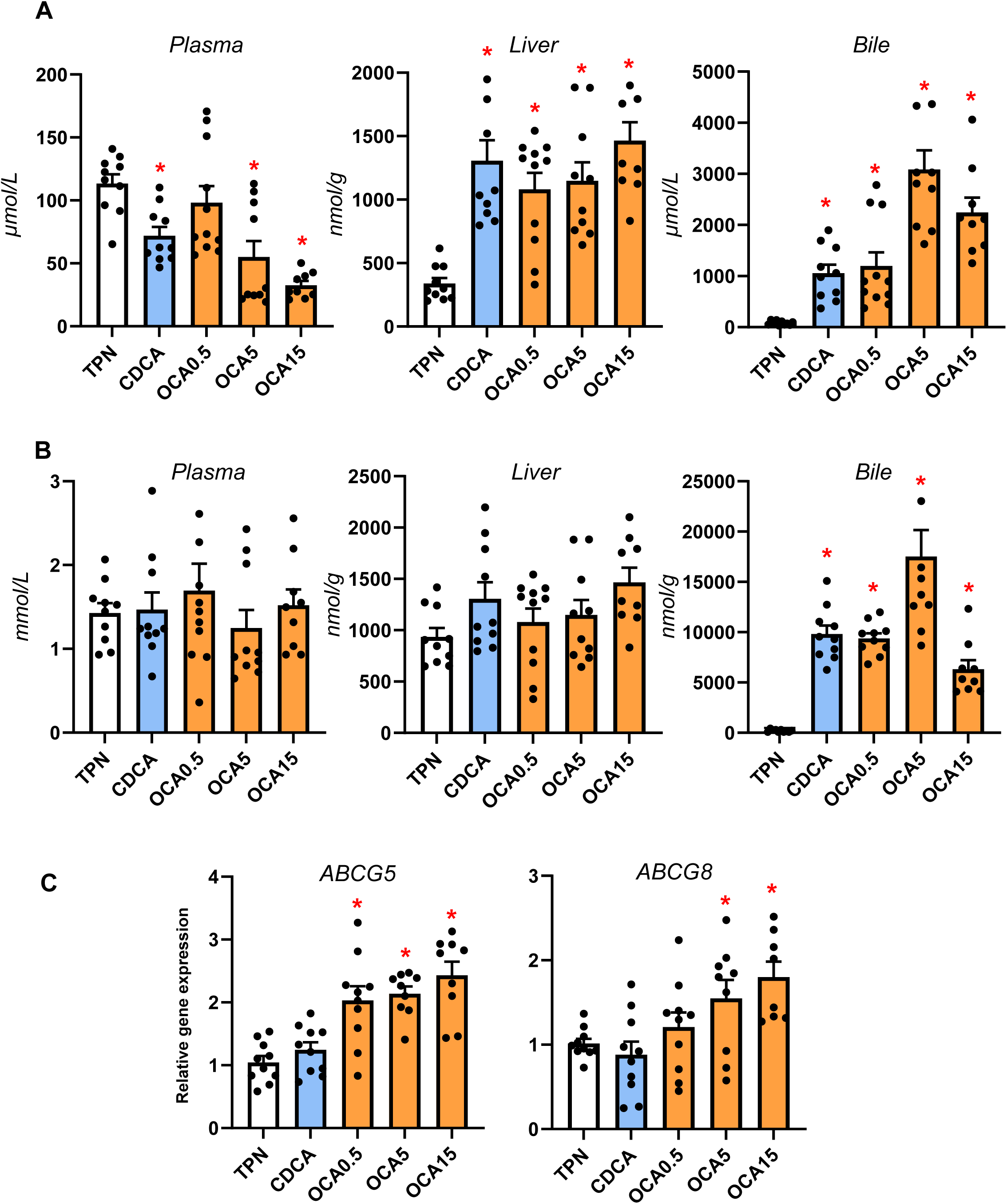
Sterol concentrations, pool sizes and hepatic transporter expression. Shown in top panel (A) are the total concentrations of phytosterols (β-sitosterol, stigmasterol, campesterol) in plasma, liver and bile in treatment groups administered TPN, TPN+CDCA (30 mg/kg/d), TPN+OCA (0.5, 5.0, 15.0 mg/kg/d). Middle panel (B) shows the concentration of cholesterol in plasma, liver and bile in various treatment groups. Bottom panel (C) shows the hepatic expression of sterol transporters, abcg5 and abcg8. Values are means with standard error, n=8-10 pigs/group. *P < 0.05 versus TPN, #P < 0.05.

### Enteral Obeticholic Acid Treatment Activates Intestinal FXR Target Gene Expression and FGF19 Secretion

We next measured the distal intestinal expression of FXR target genes given the increased flow of bile acids from the liver into the gut with enteral bile acid treatment (Figure 6). Using a porcine specific radioimmunoassay, we found a marked increase in circulating FGF19 concentration in CDCA, intermediate and high-dose OCA treated pigs. We also detected increased abundance of FGF19 protein (∼24 kDa) by Western immunoblotting in portal blood of high dose OCA-treated, but not in CDCA and TPN-treated pigs. We also observed that the intermediate and highest doses of OCA increased intestinal tissue mRNA expression of several FXR target genes, including NR0B2, FGF19, FABP6, and SLC51A.

**Figure 6.**
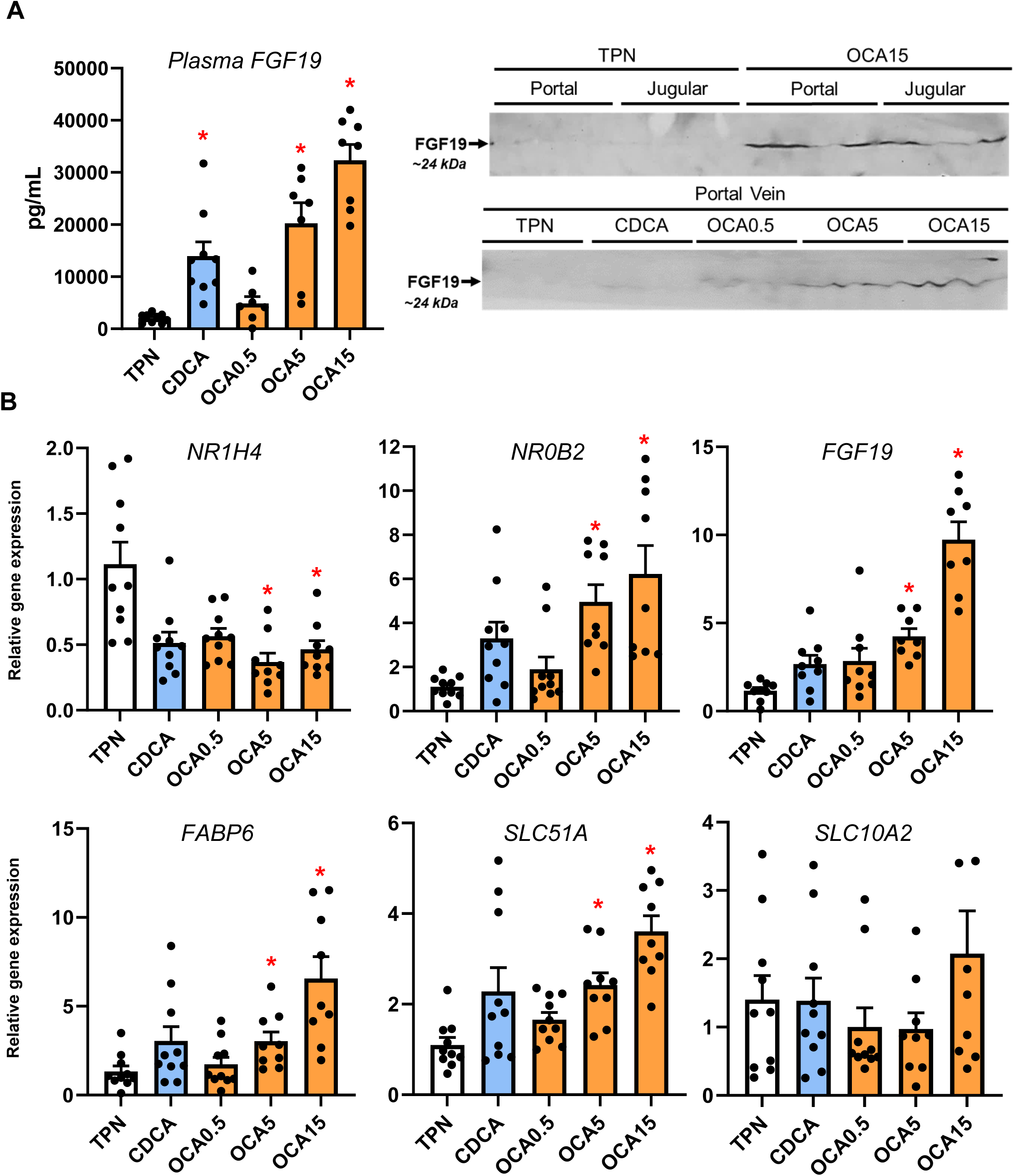
Activation of Intestinal FXR Target Gene Expression and FGF19 secretion. Shown in top panel (A) are the plasma FGF19 concentrations in treatment groups administered TPN, TPN+CDCA (30 mg/kg/d), TPN+OCA (0.5, 5.0, 15.0 mg/kg/d). Also shown are Western immunoblot images of FGF19 protein in jugular and portal venous plasma from various treatment groups. Bottom panel (B) shows the intestinal expression of FXR target genes. Values are means with standard error, n=8-10 pigs/group. *P < 0.05 versus TPN, #P < 0.05.

### Enteral Bile Acids Differentially Induce Intestinal Incretin Hormone Secretion and Mucosal Growth

Our previous studies showed that neonatal pigs given total parenteral nutrition with enteral CDCA experienced increased intestinal growth, thus we examined measures of distal intestinal growth and gut hormones (Figure 7). Our results show that CDCA and intermediate and high doses (5 and 15 mg/kg/d) of OCA increased intestinal weight, villus height and crypt cell proliferation compared to the TPN-fed group. Interestingly, we found that CDCA, but not any dose of OCA, increased the plasma concentrations of GLP-2, GLP-1 and GIP.

**Figure 7.**
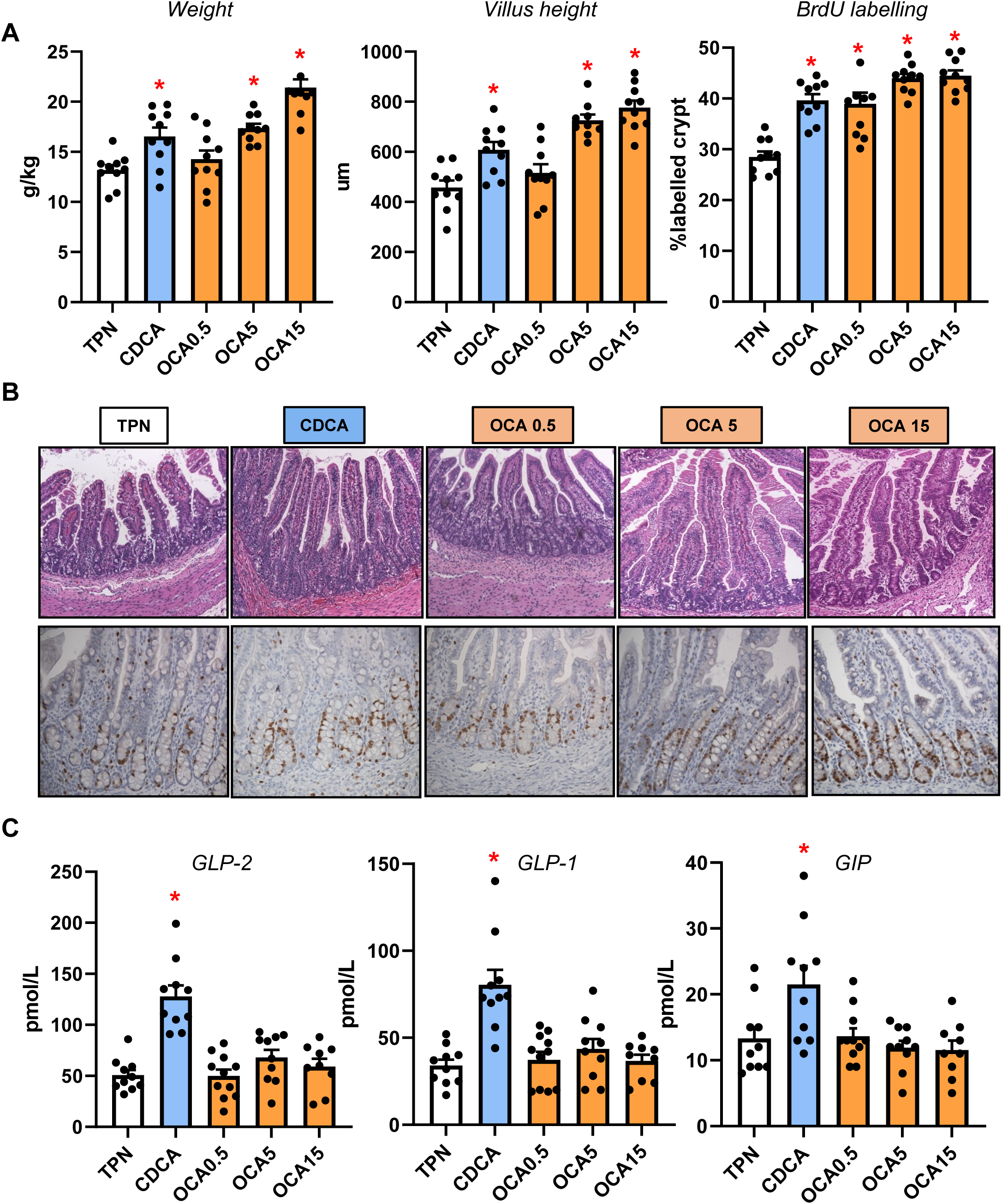
Intestinal Trophic Effects of Bile Acids and GLP-1/GLP-2 Secretion. Shown in top panel (A) are the weight, villus height and BrdU labeling in the small intestine in treatment groups administered TPN, TPN+CDCA (30 mg/kg/d), TPN+OCA (0.5, 5.0, 15.0 mg/kg/d). Middle panel (B) shows representative images of H&E stained and BrdU-stained intestinal sections from various treatment groups. Bottom panel (C) shows the plasma concentrations of gut hormones GLP-2, GLP-1 and GIP in the various treatment groups. Values are means with standard error, n=8-10 pigs/group. *P < 0.05 versus TPN, #P < 0.05.

## Discussion

Parenteral nutrition (PN) is a life-saving means of nutritional support for hospitalized infants, yet prolonged PN increases the risk for metabolic condition often termed parenteral nutrition-associated liver disease (PNALD/IFALD) that results from disruption in normal bile flow and hepatic cholestasis ^1, 2^. We previously showed in PN-fed neonatal pigs a disruption of bile acid-mediated activation of intestinal FXR-FGF19 signaling and that replacement with enteral CDCA restored FGF19 secretion and reduced serum markers of cholestasis ^14^. This previous report also showed that CDCA triggered an increase in intestinal mucosal growth and secretion of GLP-1 and GLP-2 presumably mediated by activation of enteroendocrine cell GPBAR1 receptor. Thus, because CDCA also potently activates intestinal FXR it is possible that the intestinal trophic effects are mediated via bile acid activation of FXR-dependent or independent mechanisms. In the current study, we show that a selective FXR agonist, obeticholic acid, given enterally both reduced hepatic cholestasis and induced intestinal mucosal growth, and resulted in an increase in circulating FGF19, but not GLP-1/GLP-2.

A comprehensive analysis of serum, liver and intestinal bile acid homeostasis showed that OCA dose-dependently reduced serum markers of cholestasis. This was marked by lower plasma total bile acid concentrations and increases in the gallbladder and intestinal bile acid pool sizes in OCA compared to TPN pigs. In contrast to our previous work ^14^, we did not find that CDCA reduced serum markers of cholestasis and we suspect this was because we administered a higher parenteral lipid load in current study (10 g/kg/day) compared to the previous study (5 g/kg/day). Pigs in the CDCA group showed elevated plasma and hepatic total bile acid, but also increased total bile acid content in the gallbladder and intestinal pools. There were no statistical differences in liver weight or bile duct density. These results suggest that enteral CDCA sufficiently flooded the hepatic bile acid pool, leading to spill over into the circulation while also maintaining some degree of hepatobiliary bile flow into the intestine. Yet the elevated hepatic CDCA concentrations seem to induce hepatic biliary injury marked by increased plasma direct bilirubin and GGT levels. The greater efficacy of OCA to prevent cholestasis and maintain biliary flow at lower daily doses of 5 and 15 mg/kg than CDCA (30 mg/kg/day) suggested a mechanism that mediated by its higher FXR agonist activity. Indeed, OCA is an analog of CDCA, with 100-fold higher FXR activation potency (99 nM for OCA vs. 8.66 µM for CDCA)^27^.

We next measured the bile acid profiles in the liver and distal intestinal tissue to assess the relative FXR agonist activity at key sites involved in bile acid homeostasis. We found that the concentrations of major bile acid species in pigs, namely CDCA, HCA and HDCA were as expected altered by enteral CDCA and OCA administration. In both the liver and intestine, the dominant bile acid in TPN pigs was HCA and in CDCA pigs was CDCA. Whereas in OCA groups the proportion of hepatic and intestinal OCA increased dose-dependently, such that at the highest OCA dose, OCA was the dominant species in the bile acid pool. We used literature values to calculate the FXR agonistic activity of the total bile acid pool in each tissue and as expected, estimates were highest with the two highest OCA treatments (5 and 15 mg/kg), intermediate with CDCA and lowest in the TPN pigs. A similar trend was observed in the calculated hydrophobicity of the bile acid content in the liver tissue. We also confirmed the FXR agonist capacity of the respective bile acid species and reconstituted profiles calculated from the measured liver content in each treatment group based on the induction of SLC51A expression in culture neonatal pig hepatocytes (Fig 2C). The data showed that administration of OCA at the highest doses and CDCA increased tissue FXR activation compared to control TPN group.

We then examined whether the observed tissue bile acid profiles translated into change in the expression of hepatic target genes involved in hepatic bile acid synthesis and transport. We found that hepatic cyp7a1 expression based on protein and mRNA was lower in the CDCA and all OCA groups. There was a trend for cyp4a21 and cyp8b1 to increase with the increasing OCA dose, but this was not statistically significant. We found that hepatic expression of FXR target genes NR0B2 and fgf19 were only increased at the highest OCA dose. In contrast, we found that several bile acid-regulated, ATP-binding cassette transport genes, including ABCB11, ABCC4, SLC51A, SLC10A2, and ABCB1, were dose-dependently upregulated by OCA, but not by CDCA treatment. These results suggest that the increased hepatic content of CDCA and OCA resulted in a differential effect on gene expression, where CYP7A1 was suppressed in both CDCA and OCA groups, but only the OCA treatments resulted in a robust increase in transporters involved in hepatic and hepatobiliary bile acid transport. This indicates that despite the increased tissue abundance of CDCA in CDCA-treated pigs, this was insufficient to trigger an activation of key FXR target genes involved in hepatobiliary bile acid transport. The robust activation of FXR-target genes involved in bile acid transport suggests that the higher FXR agonist activity of OCA was necessary to trigger the transporter expression that led to increased hepatobiliary bile flow into the intestine. This finding is consistent with a report in a mouse model of PNALD where parenteral treatment with an FXR agonist (GW4064) prevented hepatic injury and cholestasis and reversed the suppression of key ATP-binding cassette transporters, yet had no effect on the expression of CYP7A1 ^16^. Taken together these results imply that suppression of bile acid synthesis, vis-à-vis, reduced CYP7A1 is not sufficient to prevent cholestasis, but that concurrent FXR-mediated activation of hepatobiliary bile acid efflux into the intestine is a primary molecular mechanism.

In addition to bile acids, the hepatic accumulation of phytosterols associated with administration of soybean oil-based lipid emulsions has been linked as a molecular mechanism in PNALD. Reports in mouse models and our recent neonatal pig studies suggest that phytosterol accumulation promotes liver injury via cytokine release from activated Kupffer cells and transcriptional suppression of bile acid and sterol transporter genes, namely ABCB11 and ABCG5/ABCG8 ^7–9, 25, 28^. Our measurements of sterol concentrations suggest that both CDCA and high doses of OCA reduced plasma phytosterolemia concurrent with a marked increased concentration of phytosterols and cholesterol in hepatic bile. However, only OCA was found to induce the expression of hepatic sterol transporter genes, ABCG5/ABCG8. It is possible that the induction of FXR hepatobiliary transporters, espααecially ABCB11 and ABCG5/ABCG8, is due to the greater FXR agonism by OCA compared to CDCA that was sufficient to overcome the antagonistic effect of increased hepatocyte phytosterols concentrations. These results suggest that like, hepatic bile transporters, the upregulation of the ATP-binding cassette transporters for sterols appeared to be driven by OCA activation of hepatocellular FXR.

Finally, we examined the impact of enteral bile acid treatments on intestinal growth and bile acid signaling to assess whether the trophic actions of CDCA observed in our previous study could be mediated by FXR activation. To accomplish this, we used the selective FXR agonist, OCA, which has poor affinity for the GPBAR1 receptor, whereas CDCA has a strong capacity to activate both FXR and GPBAR1 receptors. Our results showed that both CDCA and OCA induced intestinal mucosal growth as measured by tissue weight, villus height and the crypt cell proliferation assay of BrdU labeling. Despite the stimulation of intestinal mucosal growth by both CDCA and OCA, we found that only CDCA resulted in an increase in circulating GLP-1, GLP-2 and GIP.

Interestingly, the concentrations of these key gut hormones were not different between OCA treated and control TPN pigs. We further examined the intestinal expression of FXR target genes involved in bile acid homeostasis and showed a robust dose-related increase in the expression of NR0B2, FGF19, FABP6, and SLC51A in OCA treated, but not CDCA treated pigs. We also measured a marked increase in plasma FGF19 in CDCA and the two high dose OCA groups. These results suggest that CDCA and OCA treatment, at least as the high doses, resulted in activation of intestinal FXR-FGF19 signaling that triggered increased FGF19 secretion.

The stimulation of intestinal mucosal growth by feeding CDCA or OCA enterally is notable and physiologically relevant in the context of PNALD. Considerable evidence shows that PN induces intestinal mucosal atrophy, localized inflammation and deterioration in barrier function creating a leaky intestine permeable to luminal microbial contents, such as endotoxin, that contribute to hepatic inflammation ^29, 30^. Numerous reports have shown that luminal bile acids have trophic effects on GI epithelial cell growth and cell proliferation, in the context of normal and disease conditions, such as colon cancer ^31–34^. The induction of epithelial cell proliferation by bile acids has been linked to cell mechanisms that are bile acid receptor independent involving epidermal growth factor receptor (EGFR) and mitogen-activated protein kinase (MAPK) activation ^33, 34^. The bile acid induction of epithelial cell proliferation by bile acid receptor-dependent pathways, namely FXR, suggests the loss of intestinal FXR leads to increased intestinal epithelial cell proliferation and tumor development ^35, 36^. In the current study, the increased intestinal mucosal growth induced by enteral CDCA can be explained by the activation of enteroendocrine cell GPBAR1 receptor triggering the increased secretion and circulating levels of the intestinotrophic hormone GLP-2. We also show that both CDCA and OCA increased intestinal mucosal growth and the secretion of FGF19 into the circulation. The induction of FGF19 is likely mediated by activation of intestinal FXR, but bile acid can also trigger FGF19 secretion via the pregnane X receptor (PXR)^37^. It is intriguing to speculate that the induction of intestinal epithelial FGF19 by bile acids is capable of promoting crypt cell proliferation in a paracrine or endocrine manner. Evidence suggests that intestinal epithelial cells express receptors that recognize FGF19, such as FGFR4 and β-Klotho, and cultured mouse enteroids respond to FGF19 treatment ^38^ and inhibition of FGF19 reduced growth and metastasis of colon tumor grafts ^39, 40^.

In summary, the present study showed that enteral administration of OCA to PN-fed neonatal pigs prevented cholestasis, phytosterolemia and sustained hepatic bile and sterol flow by stimulating the expression of hepatobiliary bile acid and sterol transporters. In contrast, enteral CDCA treatment was not effective in preventing cholestasis or upregulation of bile acid transporters, even though it suppressed the key bile acid synthesis gene CYP7A1. Our findings in pigs and other recent work in mice suggest that FXR-mediated activation of key hepatobiliary ATP-binding cassette transporters is sufficient to maintain hepatic bile flow and prevent liver injury. Thus, feeding a small amount of bile acids with minimal enteral feeding is a frequent clinical practice in PN-fed infants to treat cholestasis with mixed results ^41, 42^, yet the most common bile acid used is ursodeoxycholic acid, which is not an FXR agonist and has been shown to exert FXR antagonistic effects ^43^. Our results suggest that minimal enteral feeding of OCA may be an effective approach to treat PNALD in infants and thus represents a promising new indication for FXR agonists.

## Author contributions

DGB conceptualized and designed the study, conducted, and analyzed experiments and wrote the manuscript. YJ, ZF, GG, and BS designed, conducted, and analyzed experiments and edited the manuscript. SC, SL, BH, JJH, HD, JJP and II performed methods and analysis of samples and reviewed the manuscript.

---

**Supplementary Table 1.**
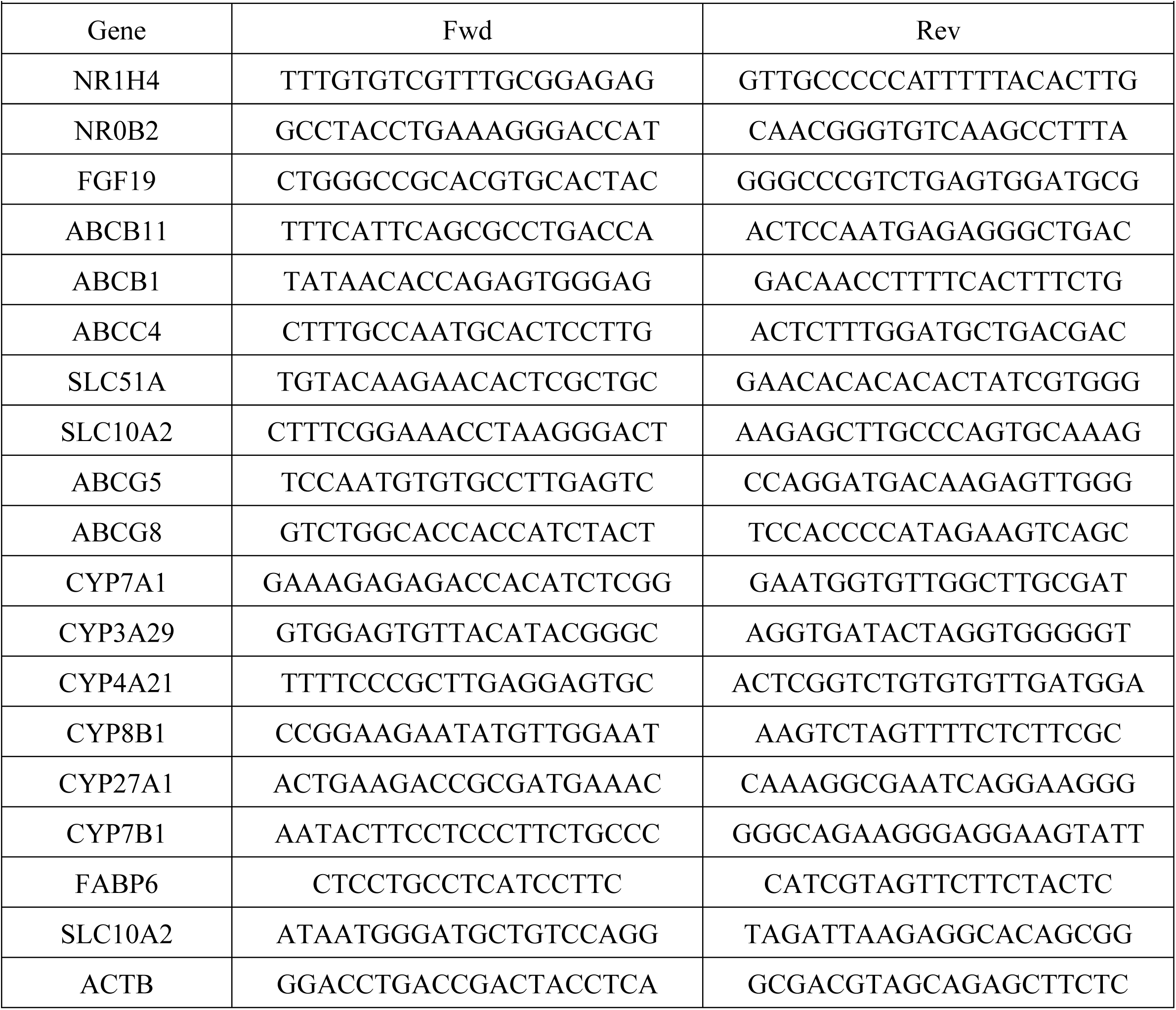
Forward and reverse primer sequences used for qRT-PCR.

